# Two routes from the Fertile Crescent led to the introduction of common vetch into Europe

**DOI:** 10.1101/2025.06.02.657542

**Authors:** Hangwei Xi, Bastien Llamas, Vy Nguyen, Mingyu Li, Qiang Zhou, Zhipeng Liu, Iain Searle

## Abstract

*Vicia sativa* (common vetch, n=6) is an annual leguminous plant with high drought tolerance and high grain protein content. Historically and still today, vetches are important for nitrogen fixation to maintain cereal yields and feeding grazing livestock. In this study, we re-sequenced the genomes of 279 common vetch accessions from a wide geographic range covering western Eurasia and North Africa to construct a comprehensive nucleotide variation map. Population structure analyses indicate that the Middle Eastern region was the centre of origin of common vetch, which then entered Europe in two distinct waves. One wave propagated by humans through Turkey into the Carpathian Basin before spreading throughout Europe via “Danubian” and “Mediterranean” routes. Demographic inferences revealed a significant bottleneck for all common vetch groups initiated during the Last Glacial Maximum, followed by an expansion that coincides with the onset of the warm Holocene epoch and the Neolithic Revolution. Interestingly, we identified selective sweeps for the flowering time regulators *VsSOC1* and *VsFTb2* in northern latitude populations suggesting that both genes are important regulators of latitudinal adaptation. Overall, we provide valuable genomic resources for conservation and breeding programs to optimize food production in the face of population growth and reduced agricultural resources.

## Introduction

The global population is projected to reach 9.7 billion people by 2050 (United Nations, 2023). Concurrently, the worldwide demand of food, particularly high-quality plant protein, is expected to undergo a disproportionate increase. Against this backdrop, *Vicia sativa*, or common vetch, emerges as a strong candidate crop to sustain future food needs. Indeed, this annual plant is a member of the Leguminosae family and is able to fix atmospheric nitrogen via symbiosis with *Rhizobia*, has a high protein content—with crude protein concentrations in its seeds averaging around 30 % (Huang et al., 2017), and is drought resilient (Tenopala et al., 2012). Together, the vetch traits position it as an attractive sustainable food source in an era of global warming and water shortage.

Vetches were identified in the archaeological record, with earliest occurrences in Syria (9200 – 8500 BCE), Turkey (6500 BCE), Bulgaria (4770 – 4728 BCE), France (4000 BCE) and Egypt (3800 BCE) (Bouby & Léa, 2006; Erskine et al., 1994; Mikić, 2016). In ancient texts Roman texts (2500 BCE), vetches were grown extensively as a “balance ration for feeding the soil and domestic animals” which we now interpret as important for providing nitrogen to the soil and a source of proteins for animals (Mikić, 2016). Vetches were valued by Roman and Greek farmers, and still today by modern farmers, for the ability to grow productively in low fertility soils and with minimal rainfall. However, a consensus on the common vetch’s locus of origin remains yet to be established. This gap in knowledge is critical given that crop breeding research often relies on a deep understanding of plants history and variation to identify traits of interest in wild variants—such as disease resistance and abiotic stress response. In the case of *V. sativa*—which is currently used as green manure or fodder for ruminant livestock—the presence of antinutritional factors within the plant, such as β-cyano-L-alanine (BCA) and γ-glutamyl-β-cyano-alanine (GBCA) (Nguyen et al., 2020), can impair nutrient absorption and pose health risks. Therefore, a better characterisation of common vetch population history and structure is required to assist processing or breeding strategies.

Whole-genome sequencing has emerged as a powerful tool to elucidate the origin, population history, and adaptation of significant plant species (Liang et al., 2019; Varshney et al., 2021; Wei et al., 2021), providing genetic information that is invaluable for breeding and conservation efforts. Yet, no largescale genetic variation dataset is available for common vetch. In this study, we sequenced a total of 308 *Vicia* accessions, including 279 globally distributed *V. sativa samples*, using Illumina short-read sequencing. We then built a comprehensive nucleotide variation dataset by aligning the sequence data to the common vetch reference genome (Xi et al., 2022), which enabled the reconstruction of the population structure and spreading of *V. sativa*, as well as the detection of selection sweeps.

## Results

### Population variation landscape of common vetch

We obtained 279 samples of *V. sativa* from seedstock centres that were globally distributed (Figure 1 a, Supplementary Table S1). Given that the archaeological records for common vetch are primarily located in West Asia and Europe (Erskine et al., 1994; Hillman, 1975), the majority of our samples were sourced from these two regions. Additionally, we gathered 29 samples of the genus *Vicia* to serve as potential outgroups for our sequent analysis. The Illumina sequencing coverage ranged from 10-30x and the average was 18.9x coverage across the 279 samples. After removing adapters and low-quality sequence data, a total of 58,008,652,420 clean reads were mapped to the 1.65 Gb *V. sativa* reference genome (n=6) (Xi et al., 2022), with an average of 99.5 % of reads primarily mapped to the reference genome (Supplementary table S1). After filtering out low-quality variant sites, we identified 118,564,208 SNPs, 7,894,289 insertions, and 9,419,947 deletions (Mean 3.46, standard deviation 10.4).

**Figure 1.**
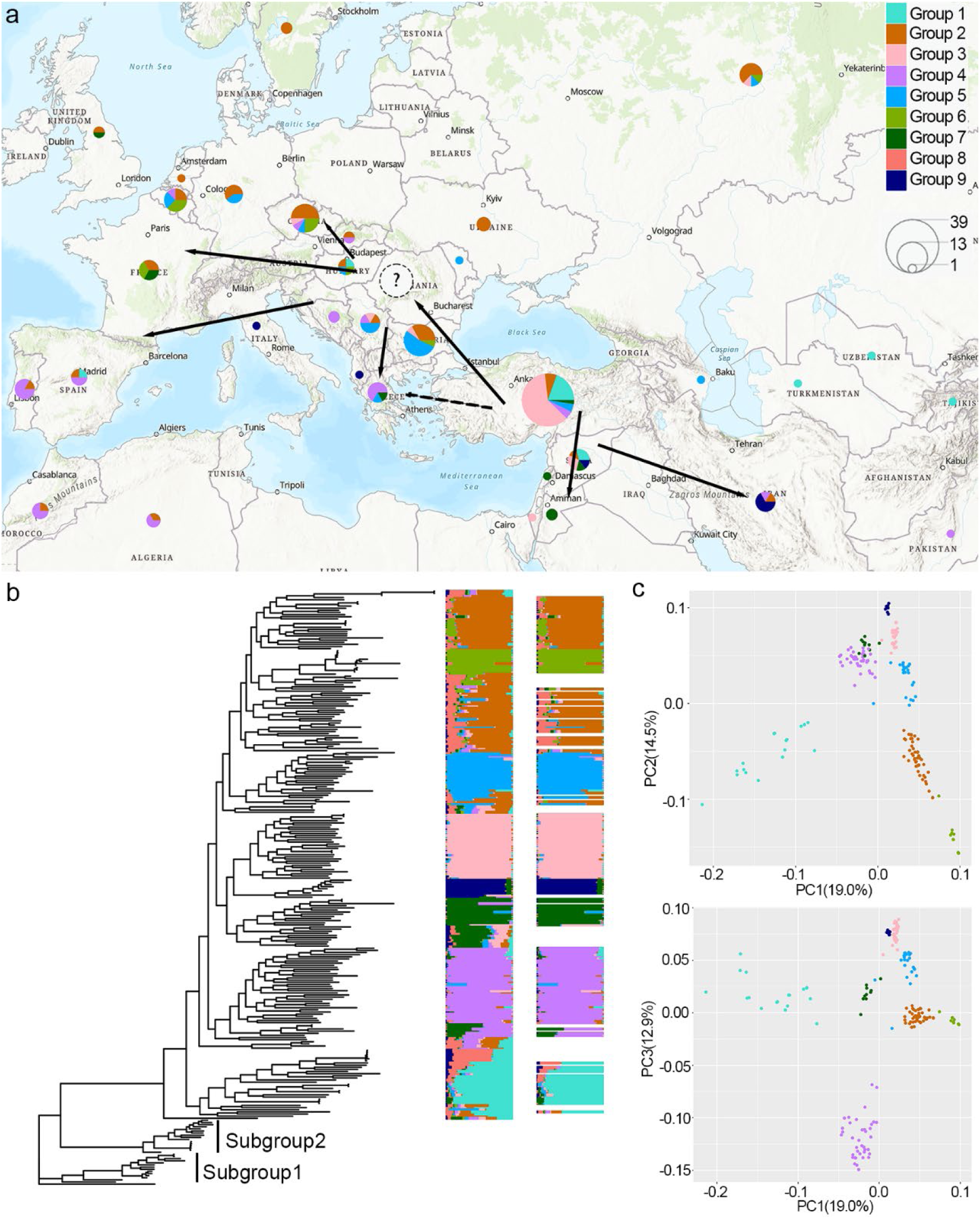
Distribution, phylogeny and population structure of common vetch. a) Geographic distribution of the 8 groups of common vetch (pie charts, size proportional to sample size) and inferred dispersal routes (arrows). The dashed arrow represents the source of a minor admixture component in the Balkan admixed population. The question mark corresponds to the ghost population in the qpGraph tree. b) Maximum likelihood tree and admixture analysis for 279 *V. sativa* samples. Subgroups 1 and 2 form the outgroup. The positions of samples in the admixture plot (right) and in the phylogenetic tree are aligned. The admixture plot shows only the samples with a major ancestry component greater than 60 %, which were kept for downstream analysis. c) Principal component analysis for *V. sativa* samples.

We observed that these nucleotide variations were uniformly distributed across the six chromosomes of *V. sativa*, with an average of one variant every 12.4 bp. When the six chromosomes were arranged in descending order of length (length shown in brackets below), we observed one variant every 12.7 bp (324.8 Mb), 12.9 bp (324.7 Mb), 12.3 bp (290.8 Mb), 12.4 bp (290.1 Mb), 11.9 bp (272.6 Mb), and 11.5 bp (148.7 Mb), respectively (Supplementary table S2). This suggests a weak trend where shorter chromosomes have a higher mutation rate. Most variants (67.3 %) were found in intergenic regions, 1.4 % in exons, and 2.8 % in introns. Within the coding regions, the ratio of non-synonymous to synonymous mutations was 1.57, which is comparable to the observed rates of 1.43 in *Arabidopsis thaliana* (Wei, 2020) and 1.37 in *Glycine max* (Yang et al., 2022).

### Population structure of common vetch

The maximum likelihood tree indicated that the IS_H04 sample was genetically closest but not included in the *V. sativa* clade (Supplementary figure S1), suggesting this specimen is not a common vetch. Indeed, we employed a K-mer analysis showed that the sequencing depth was insufficient for an accurate estimation of the genome size, suggesting that the genome size of this sample significantly exceeds that of *V. sativa*. Furthermore, only 62.6 % of reads were primary mapped to the *V. sativa* genome, further indicating it is not *V. sativa* and can thus be utilised as an outgroup for subsequent phylogenetic analyses.

After excluding sites with a minor allele frequency (MAF) less than 0.05 and a missing rate exceeding 10 %, 14,879,968 high-quality SNPs remained for subsequent population structure analysis. In the maximum likelihood tree, two basal clades exhibited substantial genetic distance from other *V. sativa* samples (Figure 1 b). The majority of samples (ten of fourteen) in the deepest basal clade, subgroup 1, were labelled as *V. sativa* subspecies nigra, providing genetic evidence for the distinction of subspecies nigra as a separate subspecies. However, nine other samples labelled as *V. sativa* subspecies nigra were scattered across other clades, indicating classification errors based on these plants’ phenotypes. For another subspecies within the sample set, *V. sativa* subspecies *amphicarpa*, we did not observe the subspecies clustering as a monophyletic group, hence not supporting its status as a distinct subspecies. Another basal clade (subgroup 2) primarily comprised samples from the Middle East and Eastern Europe, potentially representing a subspecies of *V. sativa*. A demographic history analysis revealed that these two clades diverged in their population histories from other *V. sativa* samples at least 30,000 years ago (Supplementary figure S2), earlier than the Neolithic Revolution. Therefore, these two groups are not likely to share the most recent common ancestors with the current worldwide *V. sativa* populations spread by human. We termed the remaining *V. sativa* samples, excluding subgroup 1 and subgroup 2, as the *V. sativa* core set.

The *V. sativa* core set was subjected to admixture analysis, and the lowest cross-validation error (cv-error) was obtained at K=9 (Supplementary figure S3). Samples were classified into common vetch groups 1 to 9 (referred to as groups 1 to 9 in the following text) based on the primary ancestral components identified in the admixture analysis. Interestingly, we observed widespread admixture within *V. sativa* samples. Samples containing less than 60 % of the major ancestral component were deemed admixed and were subsequently omitted from further analyses. Notably, no sample reached the 60 % threshold for the major ancestral component corresponding to group 8, and thus, no samples were assigned to this group in subsequent analysis.

Upon aligning the Admixture results with the maximum likelihood tree, there was a remarkable congruence between the groupings from the Admixture and the topological structure of the phylogenetic tree (Figure 1 b). The phylogenetic tree presents samples within the *V. sativa* core set formed a monophyletic clade, suggesting that the globally dispersed *V. sativa* originated from a singular ancestral source. Among these samples, group 1, located in the Middle East, occupied the most basal position in the tree, hinting at its close affinity with the ancestral population. This was followed by the divergence of group 4 found along the Mediterranean coast. European populations include groups 2, 6, and 5, and Middle Eastern populations include groups 3, 7, and 9. Both populations formed distinct monophyletic clades sister with each other. The topological structure of the phylogenetic tree was roughly aligned with the geographical distribution of the samples, indicating common vetch’s dispersal occurred in multiple directions.

In the PCA analysis, PC1 (19.0 %) distinctly separated group 1 from the other common vetch samples. PC2 (14.5 %) differentiated the three European groups, and correlates with an east-west gradient distribution of the *V. sativa* populations (Figure 1 c). PC3 (12.9 %) segregated group 4 from the other groups. Overall, the *V. sativa* samples exhibited a clear population structure. Despite the prominent admixture signals in numerous samples, our classification was well-suited for subsequent population genetic analyses.

### Origin and spread of common vetch

Identifying the centre of origin is crucial for crop improvement and species conservation. According to Vavilov’s centre of origin theory, the place of origin of a species typically boasts high genetic diversity (Vavilov & Dorofeev, 1992). Our findings indicated that group 1 exhibited the highest nucleotide diversity amongst all populations (Figure 2 a). Group 1 also displayed the smallest Fst and dxy values when compared to the outgroup, subgroup 1. The maximum likelihood tree constructed using IQ-TREE, the TreeMix phylogeny that accounts for gene flow, and the qpgraph based on ƒ-statistics, all showed that group 1 is the most basal amongst the *V. sativa* core set (Minh et al., 2020; Pickrell & Pritchard, 2012). This combined evidence strongly supports that *V. sativa* originated from the Middle East, where group 1 is located.

**Figure 2.**
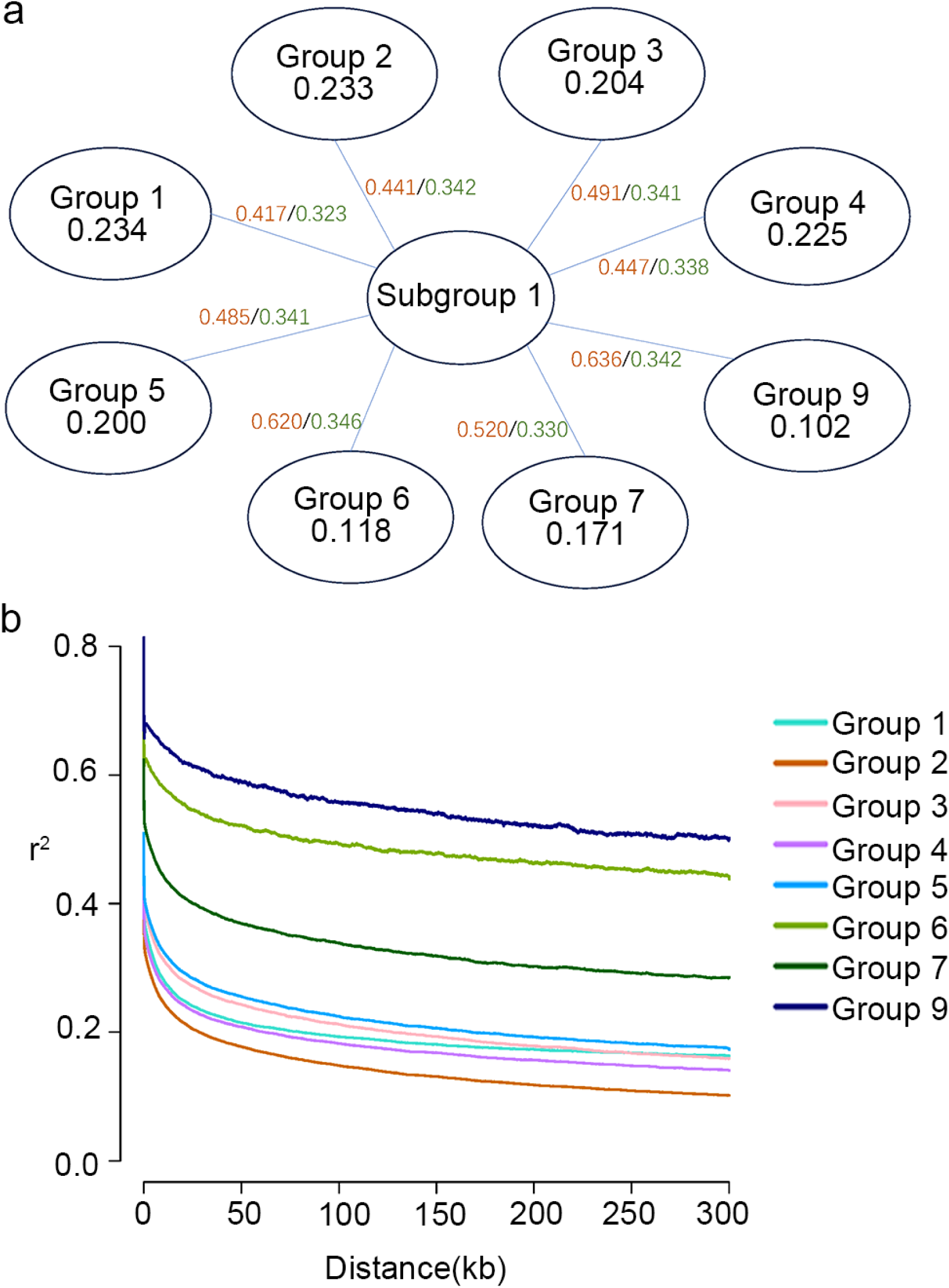
Genetic diversity and linkage disequilibrium of common vetch populations. a) Nucleotide diversity (numbers below group names in circles) of 8 *V. sativa* groups and their Fst (number in red) and dxy (number in green) compared to subgroup 1. b) LD decay of common vetch groups.

Group 2 also showed high genetic diversity. This could potentially be attributed to the group’s complex ancestral composition, as evidenced in the admixture plot (Figure 1 b), and the group’s broad geographical distribution covering eastern, western, and northern Europe. In contrast, both groups 6 and 9 had significantly reduced genetic diversity and exhibited a considerably slower decay in linkage disequilibrium (LD) compared to other groups (Figure 2 b). This could be explained by a sustained low effective population size for thousands of years (Figure 4).

*V. sativa* exhibited a complex population structure, with numerous samples showing evidence of admixture from ancestral components. This suggests frequent gene flow amongst different *V. sativa* groups. We employed TreeMix, D-statistics, and qpGraph to detect gene flow and admixture events amongst *V. sativa* groups. The TreeMix analysis strongly supported five admixture events (highest Δm in optm and lowest residual) (Supplementary figure S4, S5, S6) (Fitak, 2021). Based on qpGraph results (|Z| < 2.34), we identified two primary dispersal routes for *V. sativa* worldwide (Figure 3).

**Figure 3.**
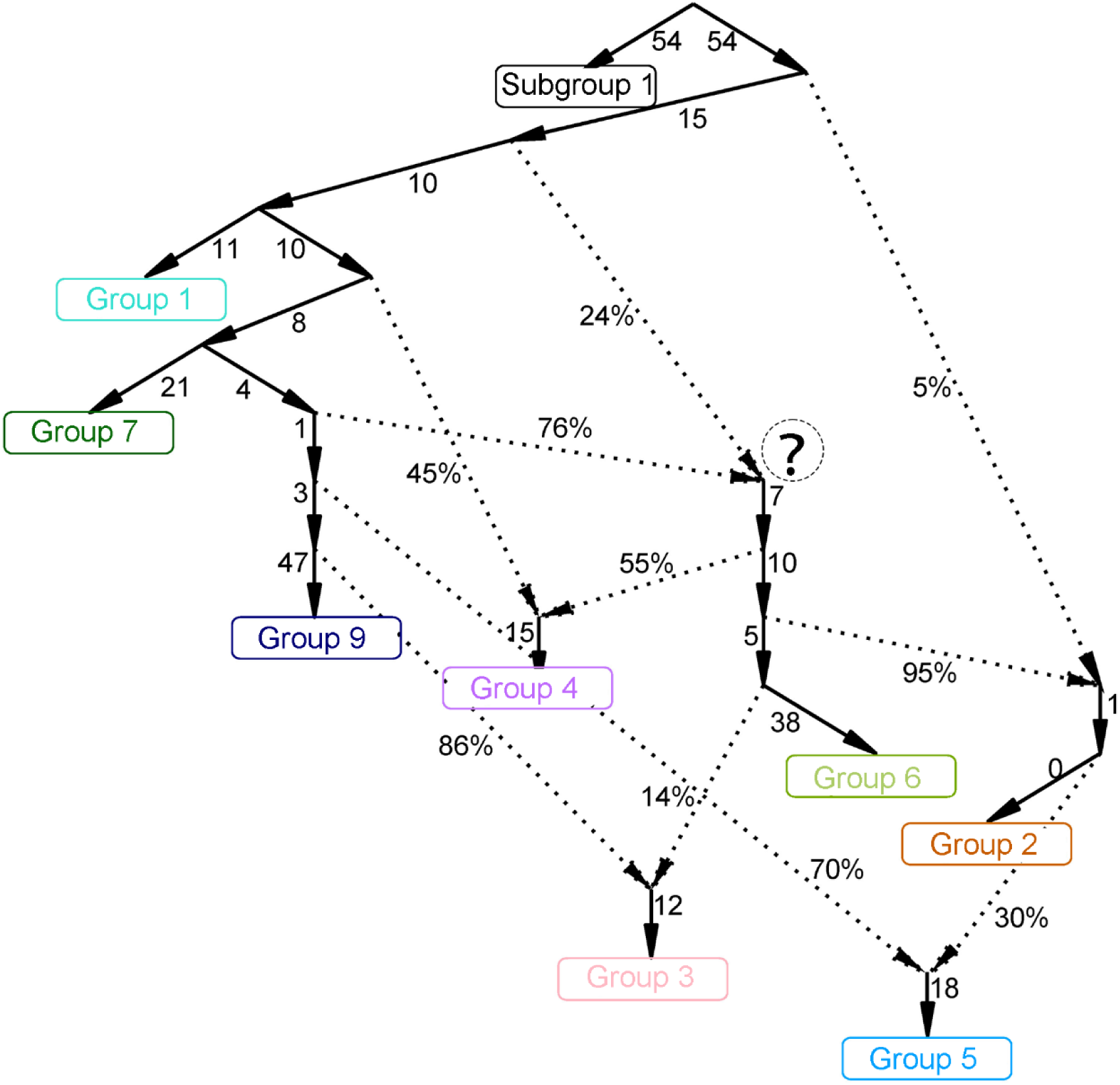
Proposed expansion route of the common vetch population. qpGraph showing relationships between common vetch groups, including Subgroup 1 as the outgroup. Numbers next to the solid arrows are for genetic drift, numbers next to the dotted arrows are for admixture proportions. The ghost population represented by the question mark is the direct ancestor of group 6, which may be the progenitor of all present-day common vetch populations distributed across Europe.

The ancestral population for the first dispersal route is group 1. From the available sampled populations, group 1 (currently in Central and southeast Asia) consistently sat in a basal position and displayed a direct phylogenetic relationship with groups 7 and 9, both located in the Middle East. This suggests that samples of group 1 initially dispersed southward, culminating in the formation of group 7 in Syria and Jordan (Figure 1 a). Concurrently, during its southern migration, a subset started moving eastward, eventually establishing group 9 on the Iranian plateau. Groups 3 and 5 are located in the Middle East and Eastern Europe, respectively, and result from admixture with their major ancestry coming from group 1. In the TreeMix tree, they form a monophyletic clade with groups 7 and 9, which indicates that the origin of groups 3, 5, 7, and 9 is common. Specifically, as group 1 was moving south, some migrated westward through Turkey to reach Eastern Europe, subsequently forming groups 3 and 5. Furthermore, both of these groups received gene flows: one from group 6 and another from the common ancestor of groups 2 and 6. Both gene flow events were validated by TreeMix (Supplementary figure S4) and D-statistics (Supplementary table S3).

The second dispersal route originated from an unsampled ghost population that resulted from an admixture event between two ancestors of groups 1 and 7 (indicated by a question mark in Figure 1 a and Figure 3). In this admixed ghost population, the progenitors of group 7 contributed the major ancestral components (76 %), while the progenitors of group 1 provided the remaining of the ancestral components (24 %).

The exact geographic distribution of this group remains uncertain because no samples were collected. This admixed group is the direct ancestor of group 2 and group 6 located in Northern and Western Europe, and it contributed 55 % of the genetic composition to group 4, which is situated along the Mediterranean coast. Considering potential dispersal routes, we speculate that this group might be situated in the Carpathian Basin region (indicated by a question mark in Figure 1 a). Subsequently, this admixed population spread to central Europe via the “Danubian” route, while the southern derivative, group 4, could rapidly diffuse to the Iberian Peninsula following a “Mediterranean” route (Marchi et al., 2022). Although the two ancestral components of this admixed group originated from distinct groups, both were traced back to the Middle East. This suggests that the admixture event could have initially occurred within the Middle East before the admixed group spread to the Carpathian Basin region. Alternatively, the ancestral components might have traversed different paths—possibly through Turkey and the northern Black Sea region—before converging and admixing in the Carpathian Basin. Genetic elements originating from the ancestral group of group 1 (the 24 % ancestral component of the mixture group in the qpGraph) spread to all common vetch populations in Europe via a secondary route. Additionally, through gene flow, these components reached the group 5 and group 3 populations located in Eastern Europe and the Middle East. These genetic elements encompass numerous alleles do not present in group 1. This explains why the LD decay rate in the European *V. sativa* populations is similar to that found in group 1, which is located at the base of the phylogenetic tree (Figure 2 b). In the qpGraph analysis, we identified a unique gene flow from the root directed into group 2 (Figure 3). This gene flow is manifested in the TreeMix tree as an edge from the outgroup (Supplementary figure S4), Subgroup1, into group 2. Consequently, this renders the LD decay of group 2 the lowest among all common vetch groups (Figure 2 b).

To investigate the demographic history of *V. sativa*, we employed SMC++ to simulate changes in the effective population size of various *V. sativa* groups across different time periods (Figure 4) (Terhorst et al., 2017). During the Last Glacial Maximum (LGM; 31–16 thousand years ago), a contraction in the effective population size of all *V. sativa* groups was observed (Figure 4). Following the conclusion of this ice age, the Middle East began to experience increased humidity, paving the way for the onset of the agricultural revolution. This is also a period when all *V. sativa* groups underwent a phase of expansion. By around 3000 BCE, this population expansion halted, and a consistent decline in the *V. sativa* population size persisted until approximately 1000 CE. This decline could potentially be associated with the widespread drought that affected Mesopotamia around 2200 BCE (DeMenocal, 2001). References to *V. sativa* began to emerge during the Roman Empire era. Literature from the 1st century BCE documented the use of common vetch as green manure and as feed for poultry and bullocks (Columela, 2020; Hernandez, 1976). Yet, during the medieval times (7th to 11th century CE), *V. sativa* remained a relatively uncommon crop in Europe (Cubero, 2018), This might correspond with a declining population history of *V. sativa* prior to the year 1000 CE. Around 1000 CE, all *V. sativa* groups, excluding group 6 and group 9, began to experience rapid expansion, which might be associated with the onset of the Medieval Warm Period (Easterbrook, 2011). In line with this, literature from the past few centuries documents that *V. sativa* spread to the New World via maritime routes and became a prevalent crop in Europe (Cubero, 2018; Weber & Schifino-Wittmann, 1999). It was also adopted as an ingredient for flour and consumed as a grain (Ramírez-Parra & De la Rosa, 2023). This mirrors the demographic expansion of *V. sativa* over the past millennium.

**Figure 4.**
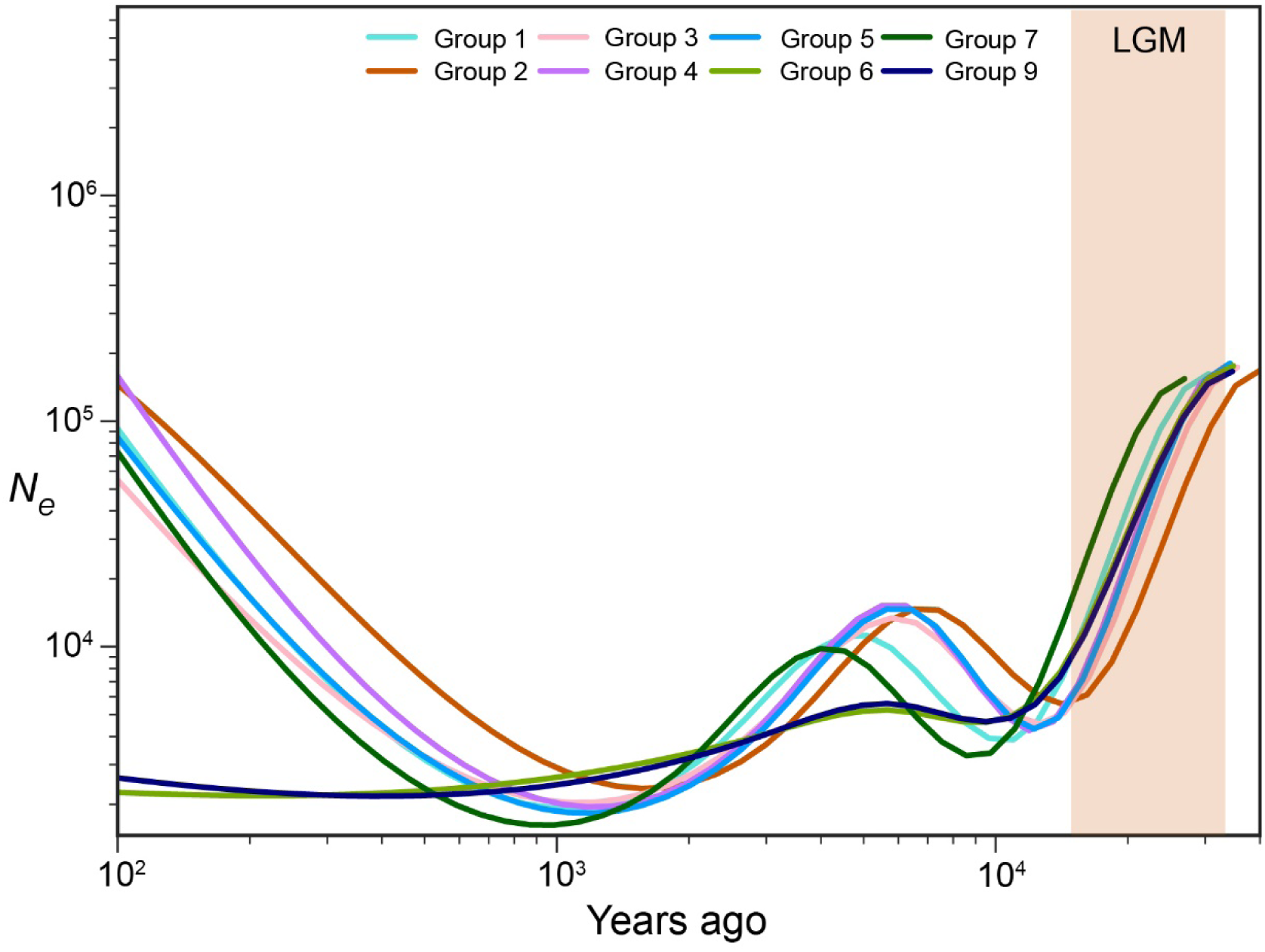
Demographic history of *V. sativa* groups from 40,000 to 100 years ago inferred by SMC++. The LGM period is represented by the light orange shaded box.

### Selection signals in common vetch genome

The population structure of *V. sativa* exhibits a notable correspondence with its geographical distribution. Various groups of *V. sativa* thrive in distinct natural habitats, having evolved over extensive periods of natural selection to become well-adapted to their local environments. Thus, they serve as ideal subjects for investigating genetic adaptation across diverse environments. Using group 1 as the ancestral group, we employed three methodologies: XP-CLR, nucleotide diversity (π) ratio, and Tajima’s D, to identify regions under selection. A threshold of 5 % was set in all analysis, defining regions detected by at least two methods as regions under selection, and those detected by all three methods as core regions under selection. In total, 153.6 Mb of regions under selection were identified across all groups, encompassing 8,532 genes. The length of the core regions under selection amounted to 42.6 Mb, containing 2,516 core genes. Following enrichment analysis of the selected genes and core selected genes in each population, we observed significant enrichment (p adjust < 0.05) in Gene Ontology (GO) terms, such as ubiquitin-like protein ligase activity, metalloexopeptidase activity, high-affinity oligopeptide transmembrane transporter activity, phospholipid transfer, and calcium ion homeostasis pathway (Supplementary table S4). In group 2, a significant enrichment of the MADS-box protein SOC1-like gene family, which plays a role in the regulation of flowering time (Lee & Lee, 2010), was detected. In the *V. sativa* genome, there are nine MADS-box protein SOC1-like genes, of which five are situated within the core regions under selection. A similar observation was noted in a study conducted on the Japanese *V. sativa* population (Shirasawa et al., 2021). We also identified an *FTb2* gene in the selected regions of group 2, which exhibits high sensitivity to long-day photoperiods and accelerated flowering time when over-expressed in *A. thaliana* (Hecht et al., 2011). In group 5, five epoxide hydrolase A genes and three allene oxide synthase 1 genes were found within the regions under selection. Both gene families are associated with jasmonic acid synthesis.

## Discussion

This study employed 279 samples of *V. sativa* primarily distributed across the Middle Eastern and European regions, along with 29 samples from the *Vicia* genus, constructing a unique and comprehensive variation map of *V. sativa*.

Phylogenetic results support the classification of *V. sativa* subspecies *nigra* as a distinct subspecies. The three most basal clades of the entire phylogenetic tree are all located in the Middle East. Even if subgroup 1 and subgroup 2 do not share the most recent common ancestor of the *V. sativa* core set, it is still anticipated that more unique genotypes could be discovered in this region.

We hypothesise that the *V. sativa* core set originated in the Middle East, entering Europe through Turkey and possibly the northern Black Sea. Subsequently, an admixed group, potentially located in the Carpathian Basin region, disseminated into Central Europe via a “Danubian” route.

All groups share a similar demographic history despite a broad geographical distribution in Western Eurasia, indicating that factors influencing vetch population size fluctuations are less likely human activity and more likely climatic shifts.

We detected adaptive signals in *V. sativa* that likely arise from the species responding to environmental changes. In group 2, SOC1-like genes and the *FTB2* gene, which were detected to be under positive selection, are associated with flowering time and photoreception, respectively. Given that group 2 primarily occupies higher latitudes with extended daylight periods, we hypothesise that positively selected alleles in these genes might delay the flowering time of *V. sativa* similar to soybean *Tof18a/SOC1a* alleles promote latitudinal adaptation of cultivated soybeans (Kou et al., 2022). This adaptation would serve to prevent frost damage to the floral organs, increasing seed set, thereby enhancing the plant’s overall fitness. In group 5, the epoxide hydrolase A genes and allene oxide synthase 1 genes are under positive selection, they are associated with jasmonic acid synthesis, which is implicated in plant defense responses under biotic or abiotic stresses (Edqvist & Farbos, 2003; Mei et al., 2006; Newman et al., 2005; Park et al., 2002; Wang et al., 2021). For instance, during herbivore or insect attacks, the synthesis of jasmonic acid surges, activating the plant’s biotic stress defences (Yan et al., 2018). This selective pressure might arise from the common vetch expansion from Turkey, which is generally more arid, to regions in Europe that are characteristically more humid, exposing it to distinct biotic and abiotic challenges. The numerous selection signals related to environmental adaptability in common vetch underscore its plasticity and indicate a significant potential for breeding improvement

In summary, this research unveils the population structure, origins, and potential dispersal routes of *V. sativa*, and identifies selection signals of environmental adaptation, offering valuable resources for the breeding and conservation of *V. sativa*. However, since the samples used in this study primarily originate from Middle Eastern and European regions, we cannot exclude the possibility that *V. sativa* may have origins in other areas. The absence of ancient DNA data also prevents us from precisely determining the temporal and spatial distributions of ancestral groups in the qpGraph. Finally, a major caveat is that no information was available to discriminate between wild and domesticated groups in our sample set, making it challenging to ascertain the exact impact of horticulture on *V. sativa*. Future research is required to address these issues.

## Material and methods

### Sample preparation and sequencing

Seeds of the 308 samples were sown in UC soil mix and grown under short-day, LED illuminated, photoperiods of 10 hours light, 22°C. Three weeks after germination, leaves from a single plant were harvested, snap frozen in liquid nitrogen and stored at −70°C until use. Leaf tissue was lysed in a Qiagen tissue lyser and DNA purified using the Qiagen Plant DNA Extraction kit. DNA concentrations were estimated by using a Qubit fluorometer. Illumina DNA libraries were prepared using the Illumina Nextera DNA Flex kit before sequencing on an Illumina NextSeq2500 (2×150bp).

### Data mapping and variant calling

Pair-end raw reads underwent trimming using Trim Galore! (v0.6.7, https://www.bioinformatics.babraham.ac.uk/projects/trim_galore) with default parameter, which was executed to excise both the adapter sequence and sequences of low-quality (Phred score < 20). The operation also discarded reads shorter than 20 base pairs. Clean reads were mapped to the *V. sativa* reference genome (Xi et al., 2022) with BWA-mem (v0.7.17) (Li & Durbin, 2009) using the default parameters. SAMtools (v1.17) (Li et al., 2009) was used to convert the mapping result into BAM format and subsequently sorted by chromosomal coordinates. Picard (v2.26.10, http://broadinstitute.github.io/picard/) was used to marking the duplicate reads in the sorted BAM file. Variant calling was performed with GATK (v4.2.5.0) (Poplin et al., 2017) in joint calling mode to enhance the sensitivity in regions with low sequencing coverage. Raw SNPs were filtered by GATK VariantFilteration with the settings: QD < 2.0 || SOR > 3.0 || FS > 60.0 || MQ < 40.0 || MQRankSum < −12.5 || ReadPosRankSum < −8.0. Given the modification to the representation of missing sites in VCF files from GATK version 4.2.3.0, BCFtools (v1.17) (Danecek et al., 2021) was used to revert the missing sites to their previous representations, ensuring compatibility with subsequent analyses. Functional annotation of the identified variants was carried out using SnpEff (v5.1) (Cingolani et al., 2012). SNPs with low MAF (<0.05) and high missing rate (>0.1) were removed to obtain a high-quality SNP set.

### Population genetic analysis

To infer the phylogenetic relationship of 279 common vetch samples, we first use SNPs situated within the coding sequence (CDS) region to construct a maximum likelihood tree among 4 *V. sativa* samples and 28 samples representing 10 species from the genus *Vicia*, by using IQ-TREE (v2.2.2.3) (Minh et al., 2020) software. Samples exhibiting the closest phylogenetic relationship to *V. sativa* group were designated as outgroup for succeeding phylogenetic tree construction. K-mer analysis was performed on outgroup samples by using jellyfish (v2.3.0) (Marcais & Kingsford, 2011). Subsequently, Orthofinder (v2.5.4)(Emms & Kelly, 2019) was used to identify the single copy ortholog between *Pisum sativum* (Kreplak et al., 2019) and *V. sativa*, and SNPs localized within these genes were used for subsequent phylogenetic tree construction, ensuring that any observed variance between the outgroup and common vetch samples originates from the identical gene. Finally, a rooted maximum likelihood tree was constructed by using IQ-TREE. The best model was determined by the model finder function, the phylogenetic tree was supported by a 1000 iteration bootstrap.

LD-filtering was performed by using Plink (v1.9) (Purcell et al., 2007) with a windows size of 50 SNPs, step length of 5 SNPs and an *r^2^* threshold of 0.5. The population structure was identified with the cluster number K ranging from 1-12 by ADMIXTURE (v1.3) (Alexander et al., 2009) on the LD purged SNPs set. Every distinct K value was run with 10 replicates and the results were consolidated by CLUMPP (v1.1.2) (Jakobsson & Rosenberg, 2007). The optimal number of the sub-population was assessed by cv-error. To address genetic component admixture, samples were tested with thresholds for retaining more than 50 %, 60 %, and 70 of the primary genetic components. It was ultimately determined that retaining samples with more than 60% of the primary genetic components achieved a balance between excessive sample removal and maintaining noticeable admixture. The ggtree (v3.8.0) (Yu et al., 2017) package was used to align the ADMIXTURE result to the phylogenetic tree by sample names. PCA was performed on the same dataset using Plink, the first 10 principal components were calculated, wherein the first 3 principal components were plotted. LD between each pair of SNPs within each population was calculated using PopLDdecay (v3.4.2) (Zhang et al., 2019) with the default parameter. Pixy (v1.2.7) (Korunes & Samuk, 2021) was used to calculate genetic differentiation (Fst) and absolute divergence (dxy) between each group, as well as the π within each group with a 10kb nonoverlapping window.

### Demographic history of common vetch

The effective population size was estimated using SMC++ (v1.15.2) (Terhorst et al., 2017). A subset of ten random samples was selected from each group, except for group 9, which only contained seven samples. According to the quality standard of GATK VariantFilteration, the coordinates of low-quality SNPs were masked in this analysis. We used a mutation rate of 8.02 × 10^-9^ per site per year (Xi et al., 2022), and a generation time of one year. SMC++ split was used to infer the split times between different populations. However, due to the significant gene flow between the populations in this study, the inferred split times could be highly biased using this method, so they are not discussed further.

To detect the genetic introgression among common vetch groups, we first use Treemix (v1.13) (Pickrell & Pritchard, 2012) to construct a ML tree, varying the migration edge quantity between 1 and 8. For each specific number of migration edges, 20 replicates were conducted utilizing unique random seed, and the phylogenetic tree was rooted by designating subgroup 1 as an outgroup. Following this, OPTM (v0.1.6) (Fitak, 2021) package was used to infer the most optimal count of migration edges. Subsequently, we used qpgraph, a component of the ADMIXTOOLS2 (v2.0.0) (Maier et al., 2023) package, to model the interrelationships and admixture amongst the groups, once again treating subgroup 1 as the outgroup. D-statistics were calculated using the identical package and outgroup.

### Selective sweep detection

To detect intra-population selection signals, we applied three distinct methods. First, we calculated the π ratio for each group, specifically dividing the π value of the target population by the mean π values of all other remaining groups. Next, we utilized VCFtools (v0.1.16) (Danecek et al., 2011) to compute Tajima’s D for each group. Lastly, a XP-CLR test was performed using XP-CLR (v1.1.2) (Chen et al., 2010) software, group 1 was used as the reference group. The recombination rate was calculated by dividing the total genetic distance of *V. faba* (Ellwood et al., 2008) by the genome size of *V. sativa*. All three analysis are using a 10kb nonoverlapping window. In the three analyses conducted, regions falling within the top 5% in any two of the analysis were considered as the region under positive selection. Subsequently, we extracted genes present in these positive selected regions and conducted an enrichment analysis of these candidate selected genes using the ClusterProfiler (v4.8.1) (Wu et al., 2021) package.

## Data Availability

Sequencing data is available at NCBI BioProject PRJNA1218287.

## Acknowledgements

This research was supported by grants from the the Australian Research Council (ARC, LP200200957), Leading Scientist Project of Gansu Province (23ZDKA013), the Gansu Provincial Science and Technology Major Projects (22ZD6NA007).

