## Supplementary figures and images for "Two routes from the Fertile Crescent led to the introduction of common vetch into Europe"

### Supplemental Figure 1

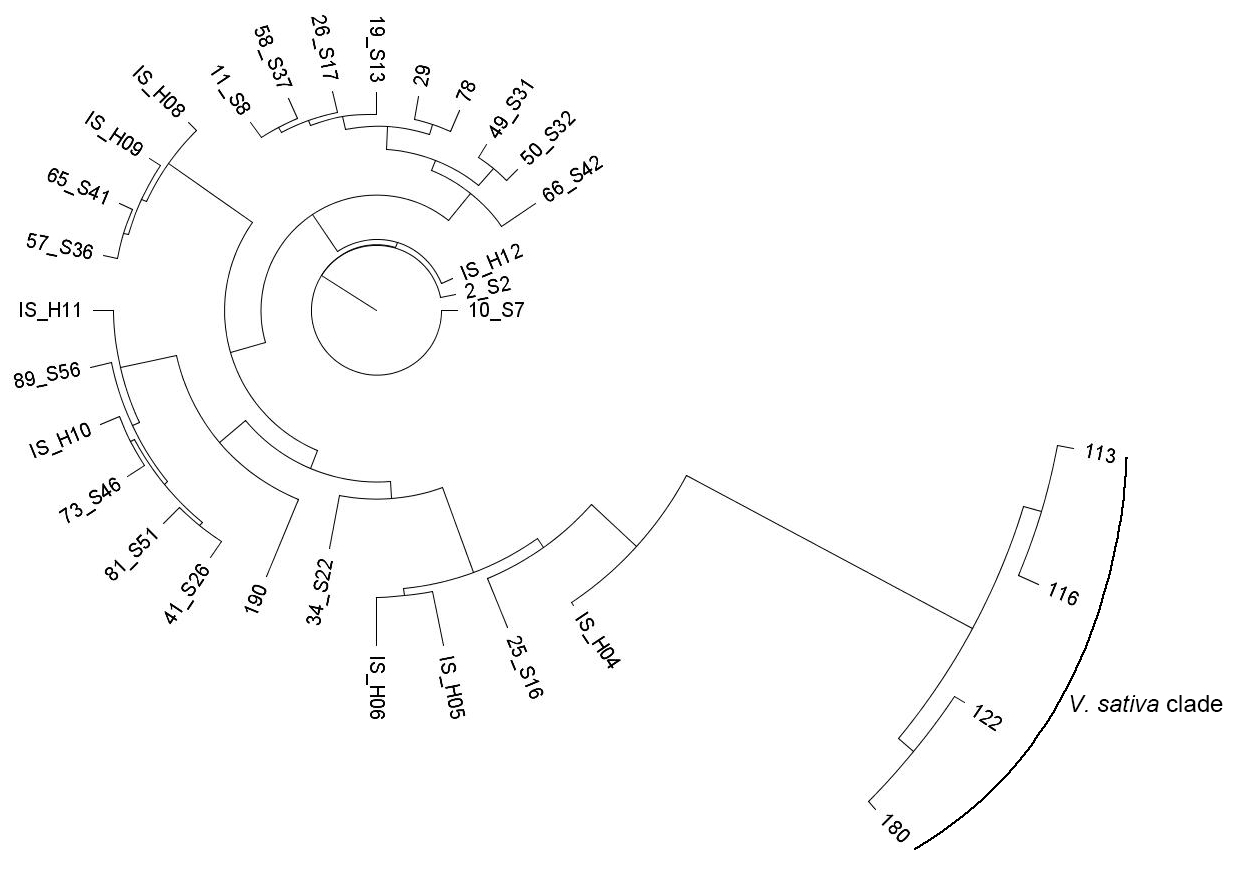

### Supplemental Figure 2

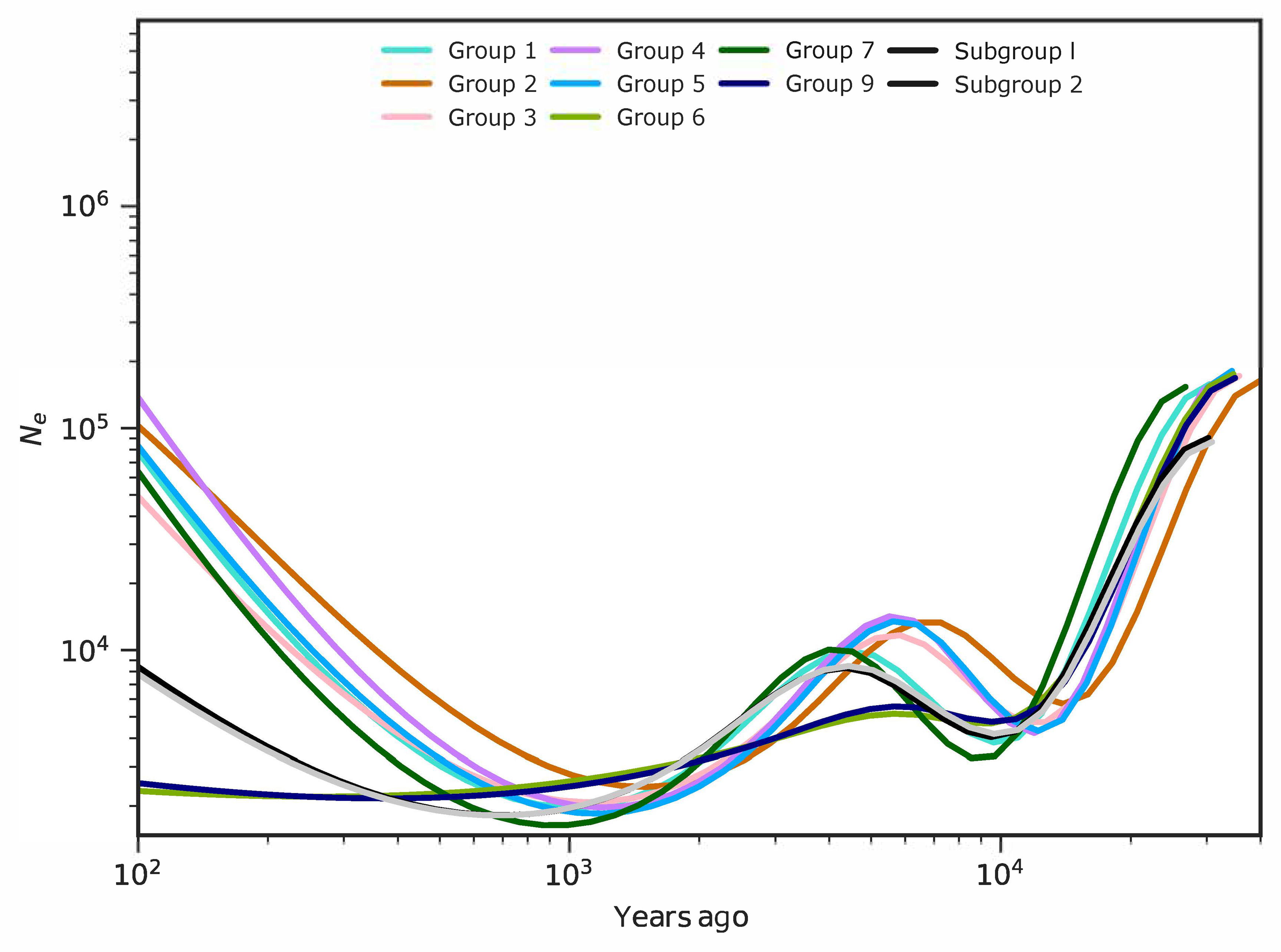

### Supplemental Figure 3

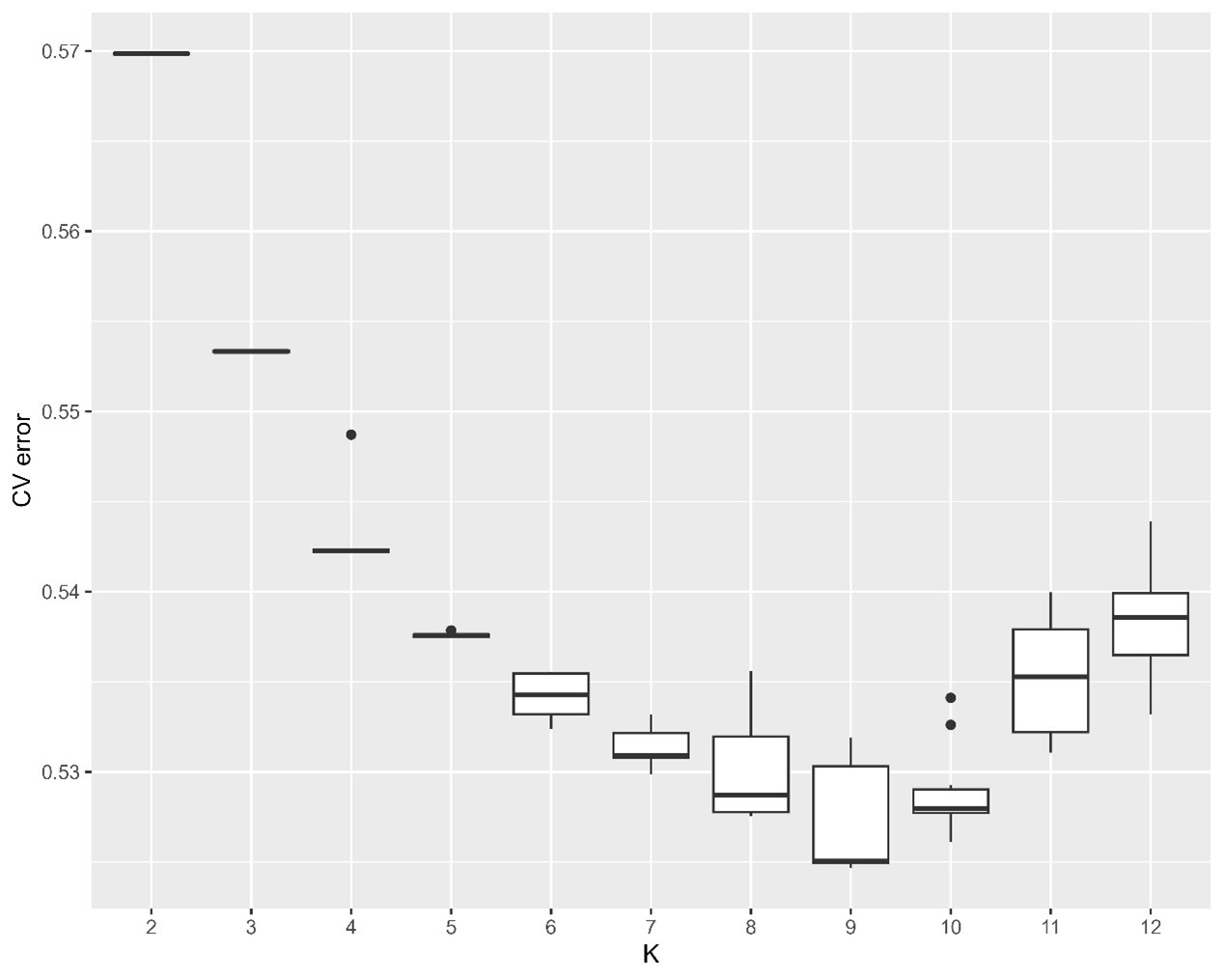

### Supplemental Figure 4

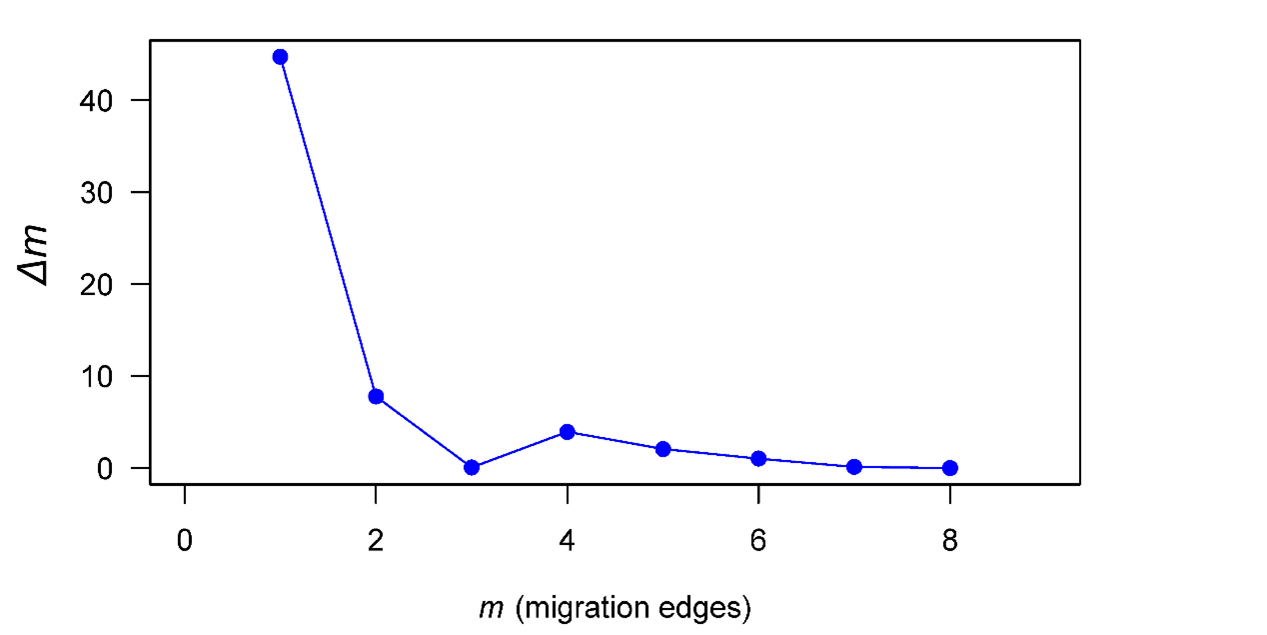

### Supplemental Figure 5

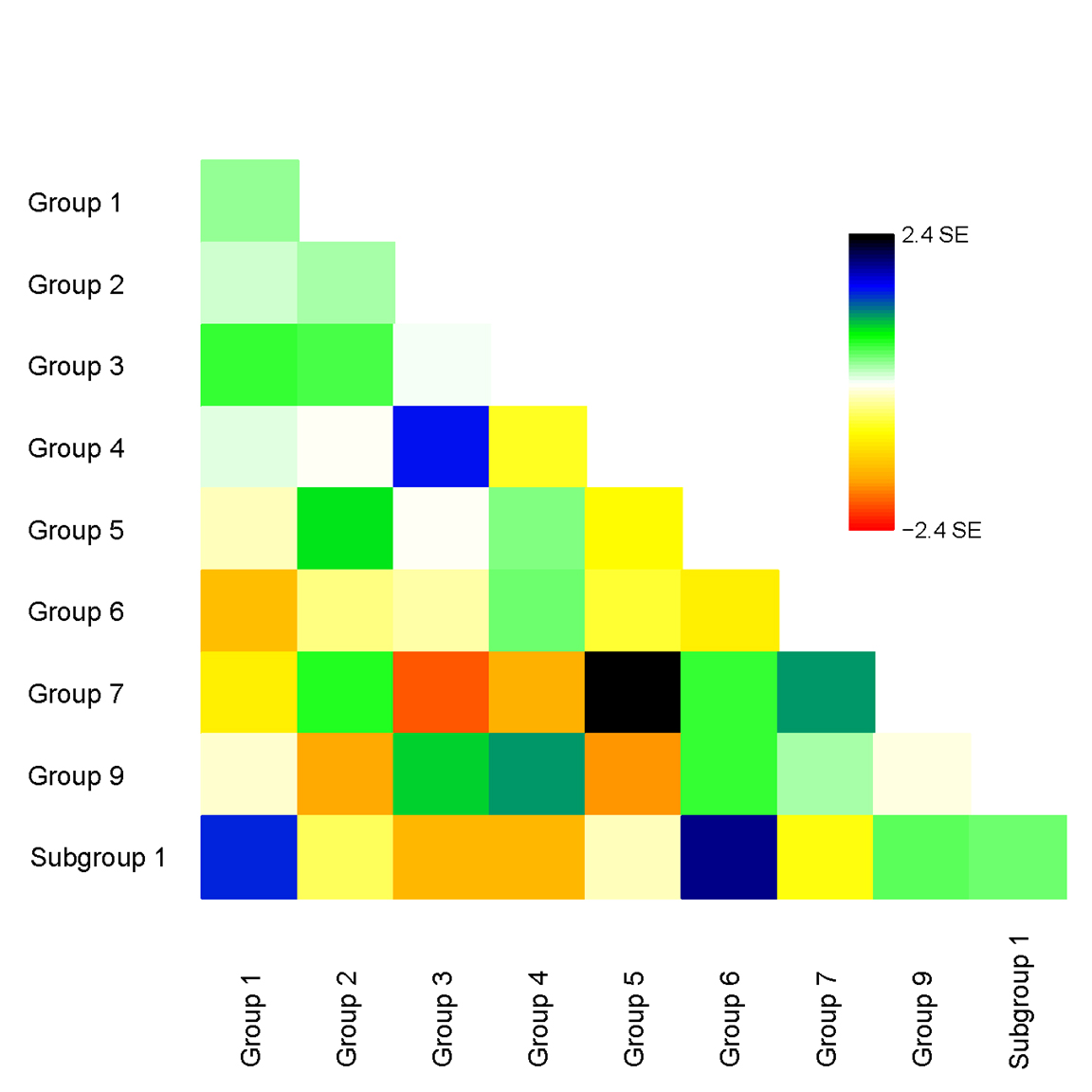
